## Supplementary Figures for "A ubiquitous GC content signature underlies multimodal mRNA regulation by DDX3X"

### Supplementary Figures – Jowar et al, 2023

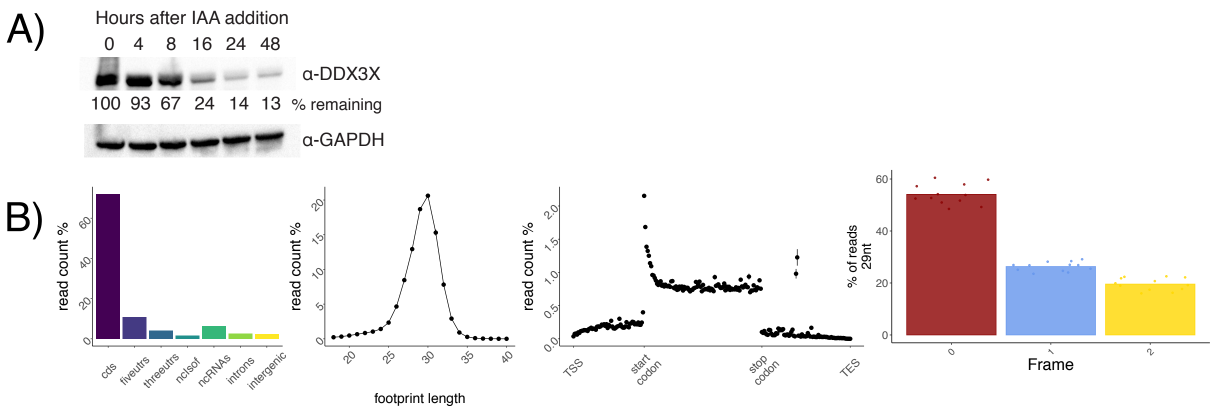


**Supplementary Figure 1**. Degron and Ribo-seq quality control

A) Immunoblot indicating DDX3X and GAPDH levels during the IAA time course. Percent DDX3X is calculated as the DDX3 intensity normalized to GAPDH across two replicates. B) Read mapping locations are shown on the far left, followed by footprint lengths distributions, a metatranscript coverage plot, and frame resolution analysis.

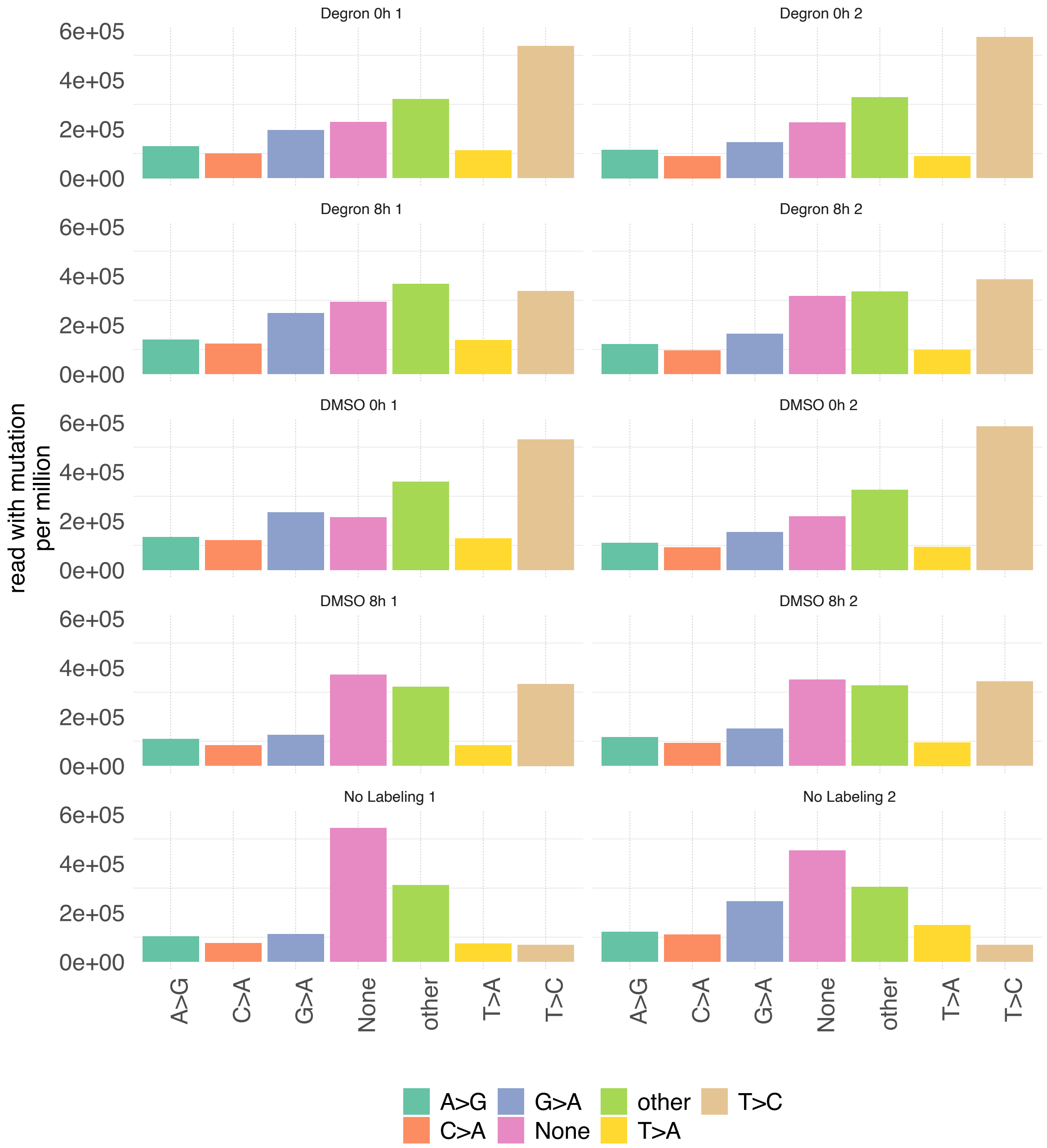


**Supplementary Figure 2**. SLAM-seq quality control

Read counts harboring different mutations are shown across datasets. The SLAM-seq dataset contained background sequencing or RT errors, leading to elevated mutations other than T>C, even without 4sU labeling. However, a decrease in T>C harboring reads can be observed (indicating mRNA decay) for the 8h time points.

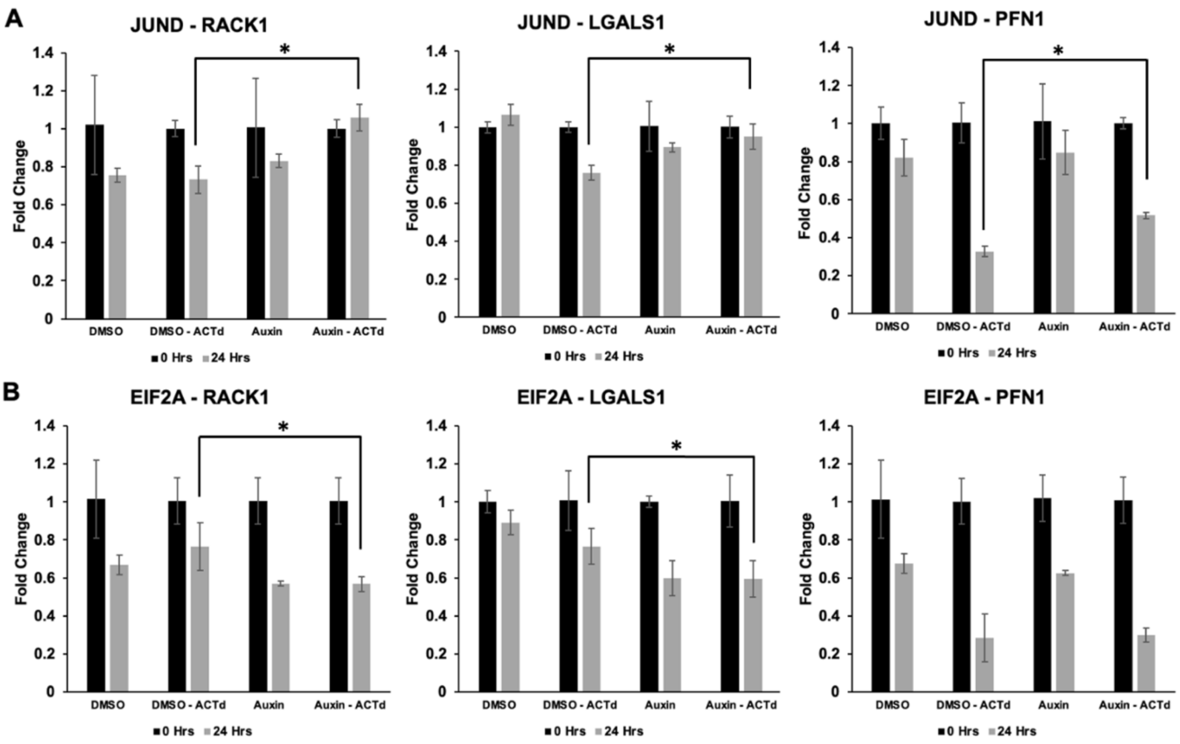


**Supplementary Figure 3**. qPCR validation of mRNA stability changes

JUND and EIF2A stability changes via qPCR. A. Histogram representing JUND expression by RT-PCR via taqman probes. B. Histogram representing EIF2a expression by RT-PCR via taqman probes. Fold change was normalized to RACK1, LGALS1, or PFN1. *p-value < 0.05. Performed in triplicate.

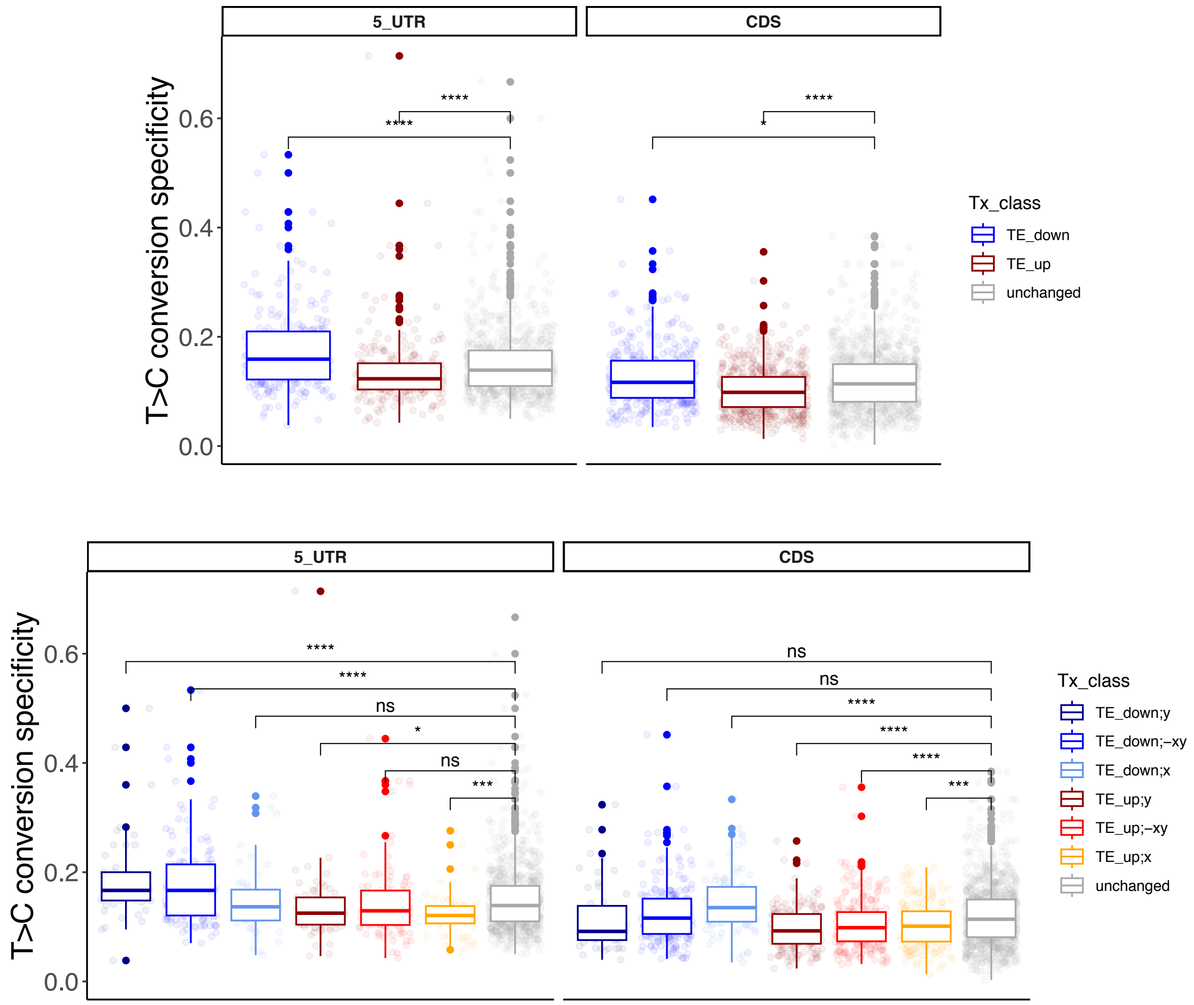


**Supplementary Figure 4**. mRNA binding pattern on different regulated mRNAs.

T>C conversion specificity on the y-axis (as defined in Calviello and Venkataramanan et al, NAR 2021) is plotted against different regulated transcripts, for both 5'UTR and CDS peaks. Significance values come from two-sided Wilcoxon test against the control gene group.

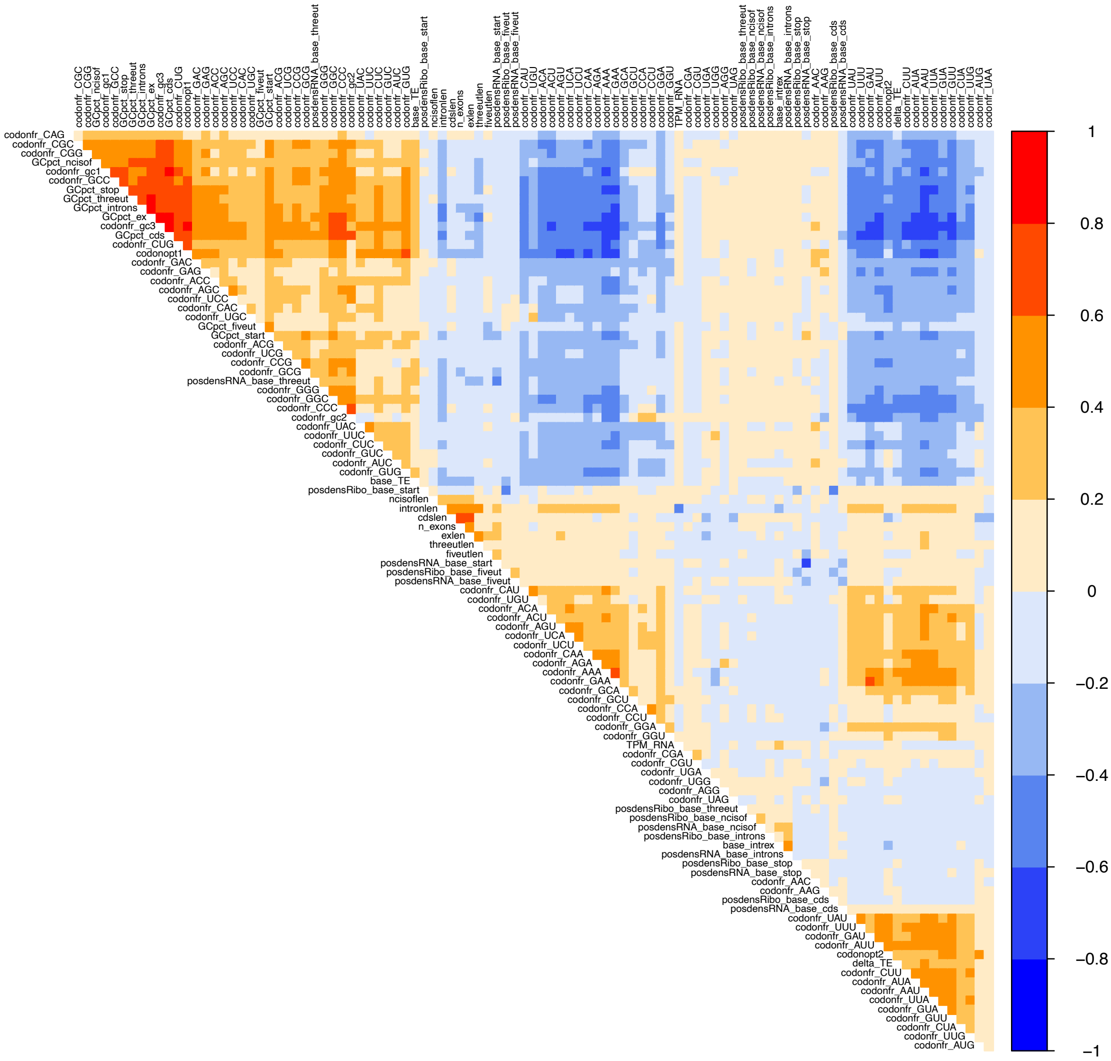


**Supplementary Figure 5**. Correlation among different mRNA features

Pearson correlation values across the features used as input for the Random Forest and Lasso regression models. As expected, many features (e.g. GC content along transcripts regions and GC-rich codon frequencies) show high level of correlation.

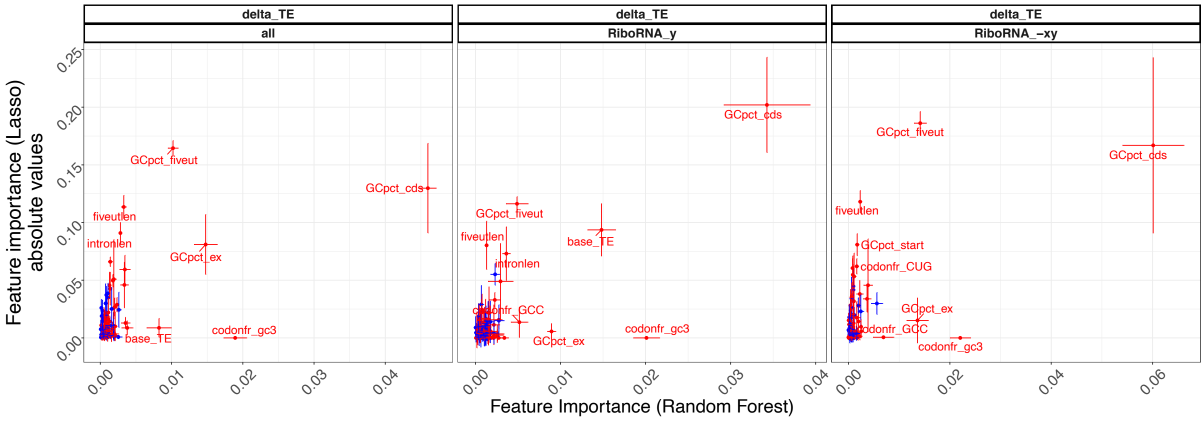


**Supplementary Figure 6**. Comparison between Lasso and Random Forest feature selection results

Feature importance according to the Random forest on the x axis, with Lasso coefficients (taking the absolute value) on the y axis. Error bars calculated from 5-fold cross-validation estimates. In red the most relevant features.

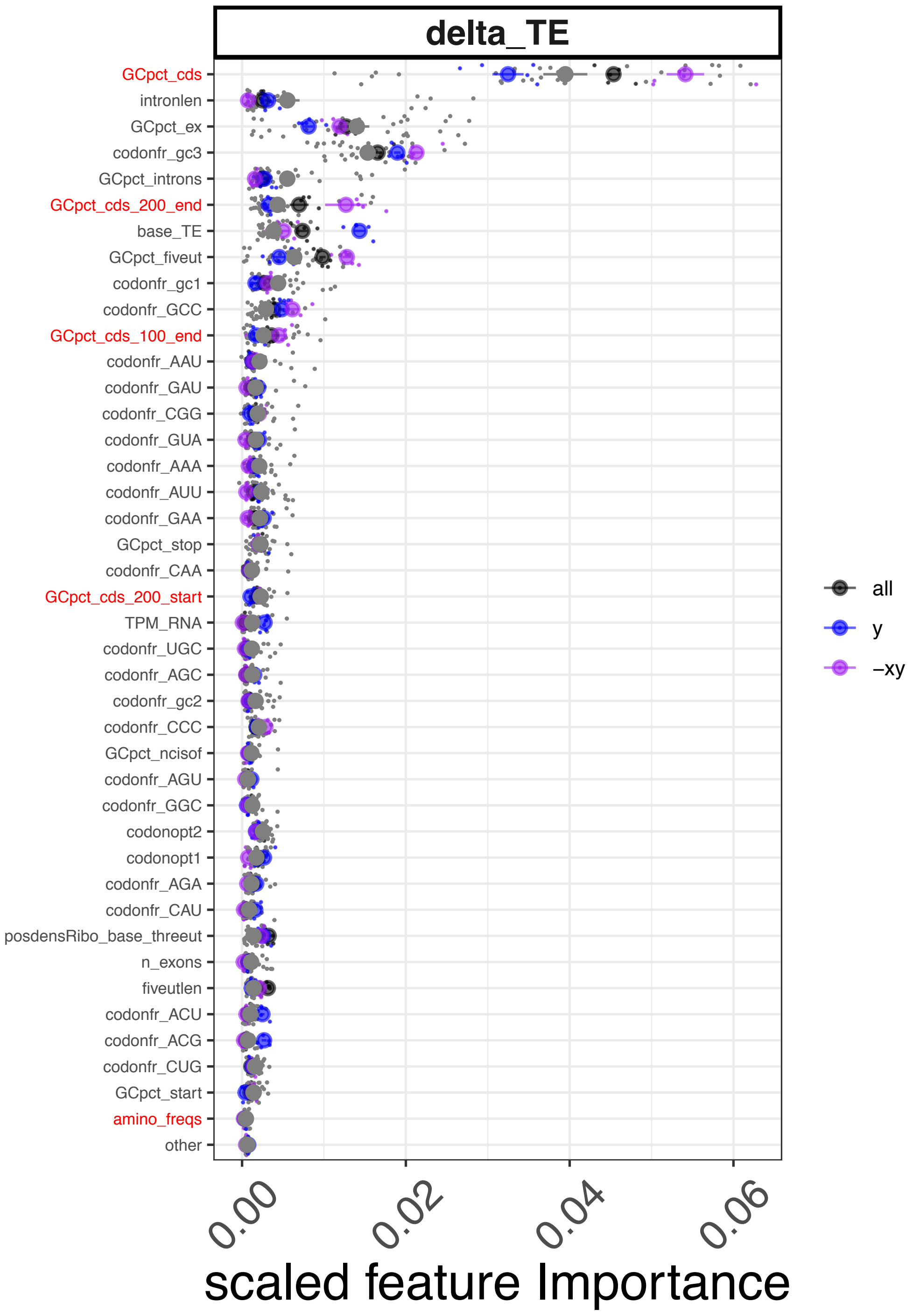


**Supplementary Figure 7**. Importance plot with additional features

Predictive power of different features in quantifying translation regulation, with their importance values plotted on the x axis. New variables, alongside GCpct_cds, are colored in red.

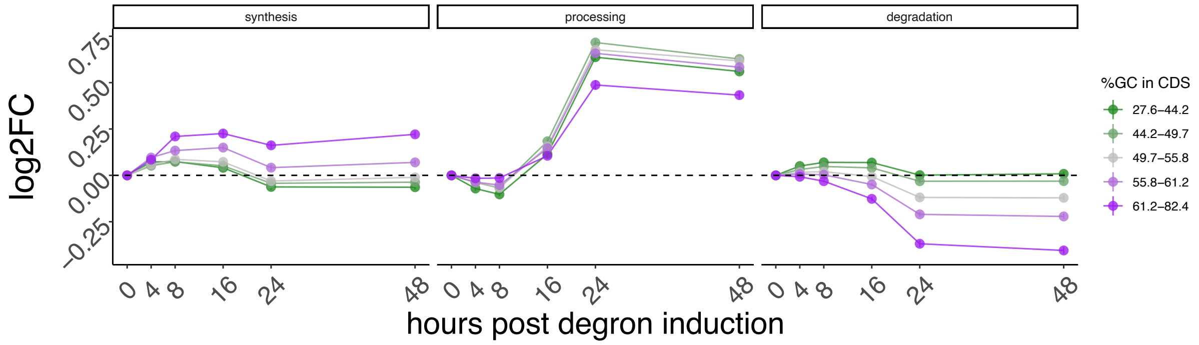


**Supplementary Figure 8**. mRNA dynamics divided by GCcds values.

Synthesis, processing and degradation values, as inferred by INSPEcT, partitioned by GCcds values.

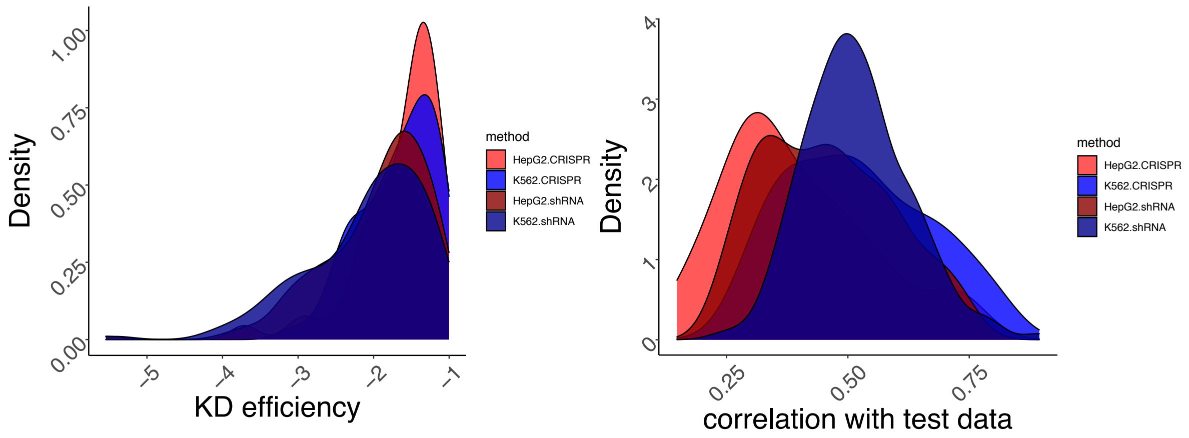


**Supplementary Figure 9**. ENCODE RBP data analysis overview.

RBP knockdown efficiency (log2FC) across methodologies in the ENCODE dataset (on the left). On the right, performance of the Random Forest model across ENCODE knockdown methods and cell lines.

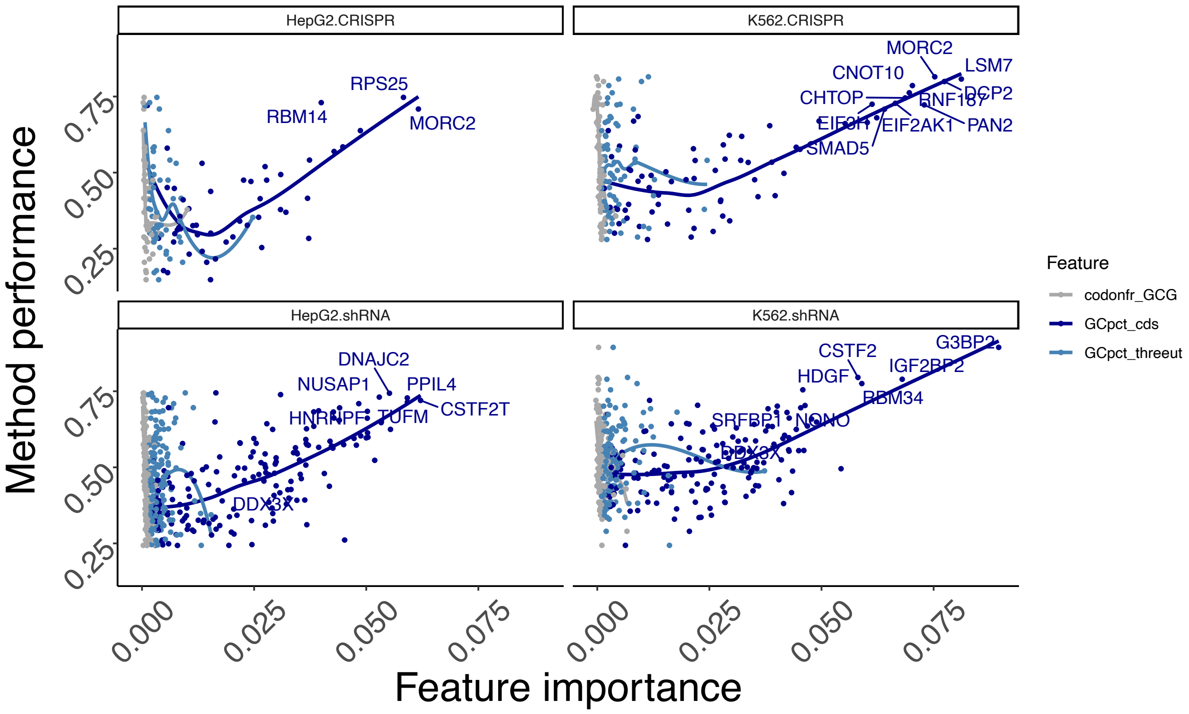


**Supplementary Figure 10**. *GCcds* importance across the entire ENCODE dataset

Figure R3: Model performance (spearman correlation between predicted and real values on unseen test data) on the y axis, with importance of 3 example features variables (indicating their predictive value) on the x axis. Top knockdown experiments, together with DDX3X, are show with labels. Data shown separately for each ENCODE knockdown strategy. The linear relationship between *GCcds* importance and model performance indicates its relevance as the top predictor of RNA changes in dozens of datasets.

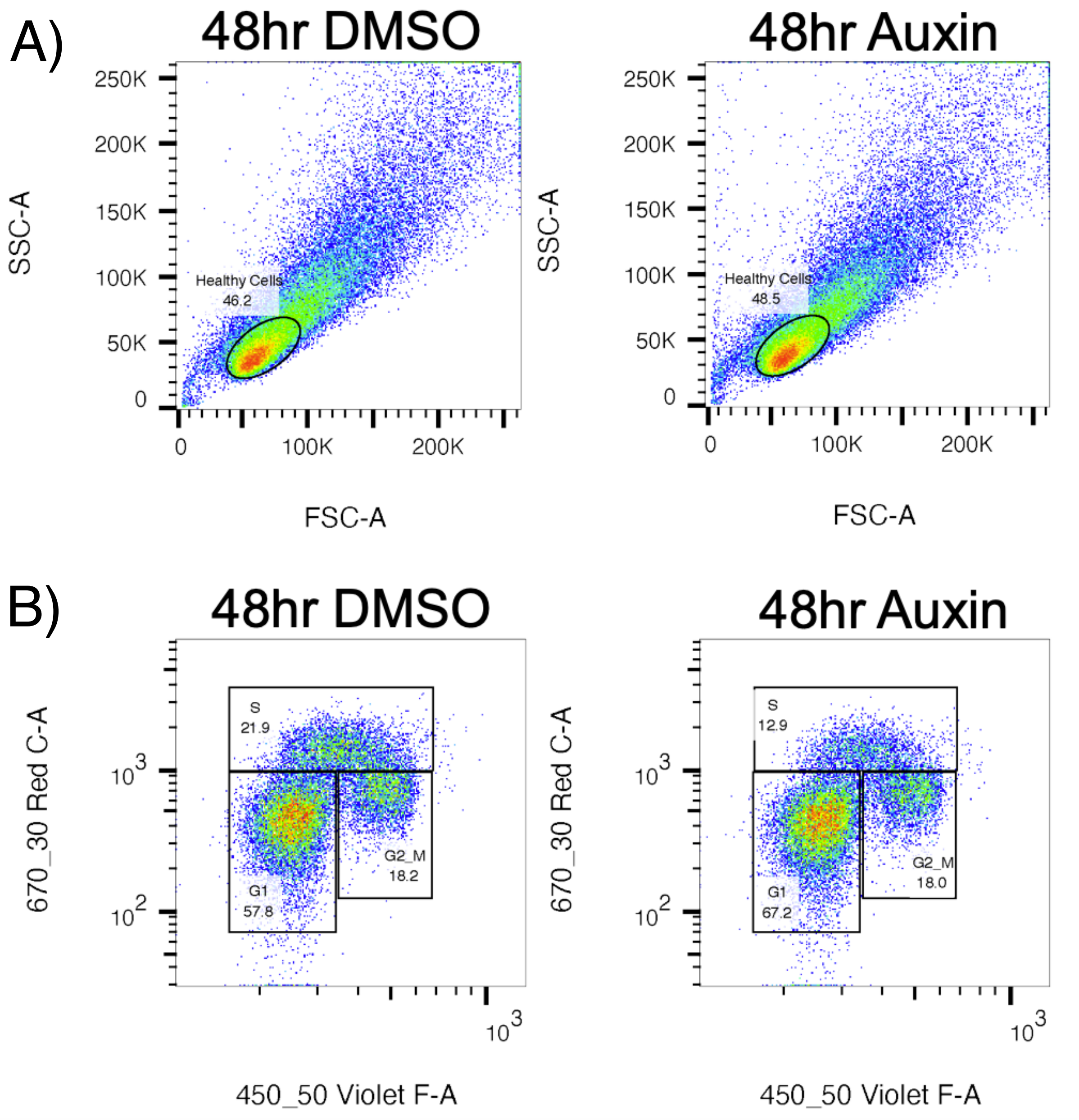


**Supplementary Figure 11**. Cell cycle staging analysis.

Cell-cycle analysis of HCT116-DDX3-degron cells treated with either DMSO or Auxin (500uM) for 48 hrs. 1 representative replicate of 3 total replicates is shown. A) Single healthy cells were gated on their Forward (FSC-A) and Side Scatter (SSC-A), B) Cells currently undergoing DNA synthesis incorporate EdU (670-30 Red C-A Channel) and FxCycle Violet Stain (450-50 Violet F-A Channel) labels overall DNA content. Cells can then be separated into those in G1, S, and G2-M phases of the cell cycle.

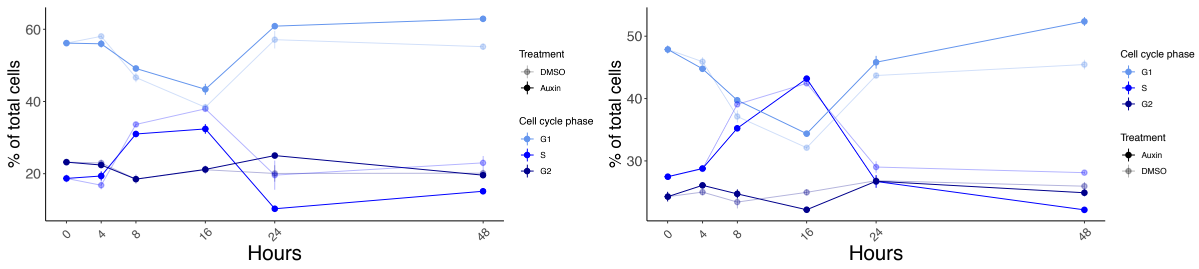


**Supplementary Figure 12**. Cell cycle dynamics after DDX3X degradation.

Percentage of cells in different stages of the cell cycle along the degron time course. Two independent experiments are shown. Values for degron and control are shown with different transparency values.

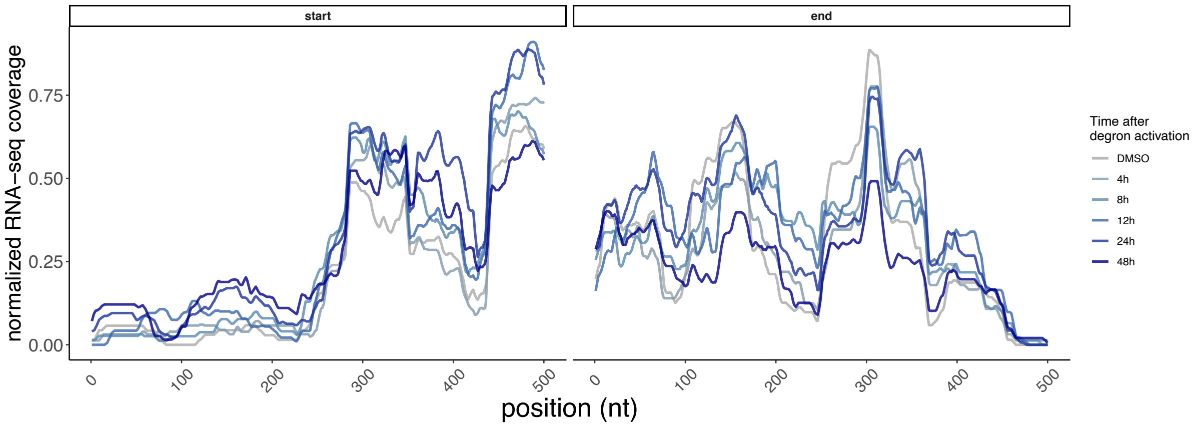


**Supplementary Figure 13**. Example of RNA-seq coverage changes across the DDX3X degron time course.

RNA-seq coverage tracks around 5’ and 3’ ends of the stabilized mRNA from the *CSRNP2* gene. Coverage values were 0-1 normalized for each dataset.

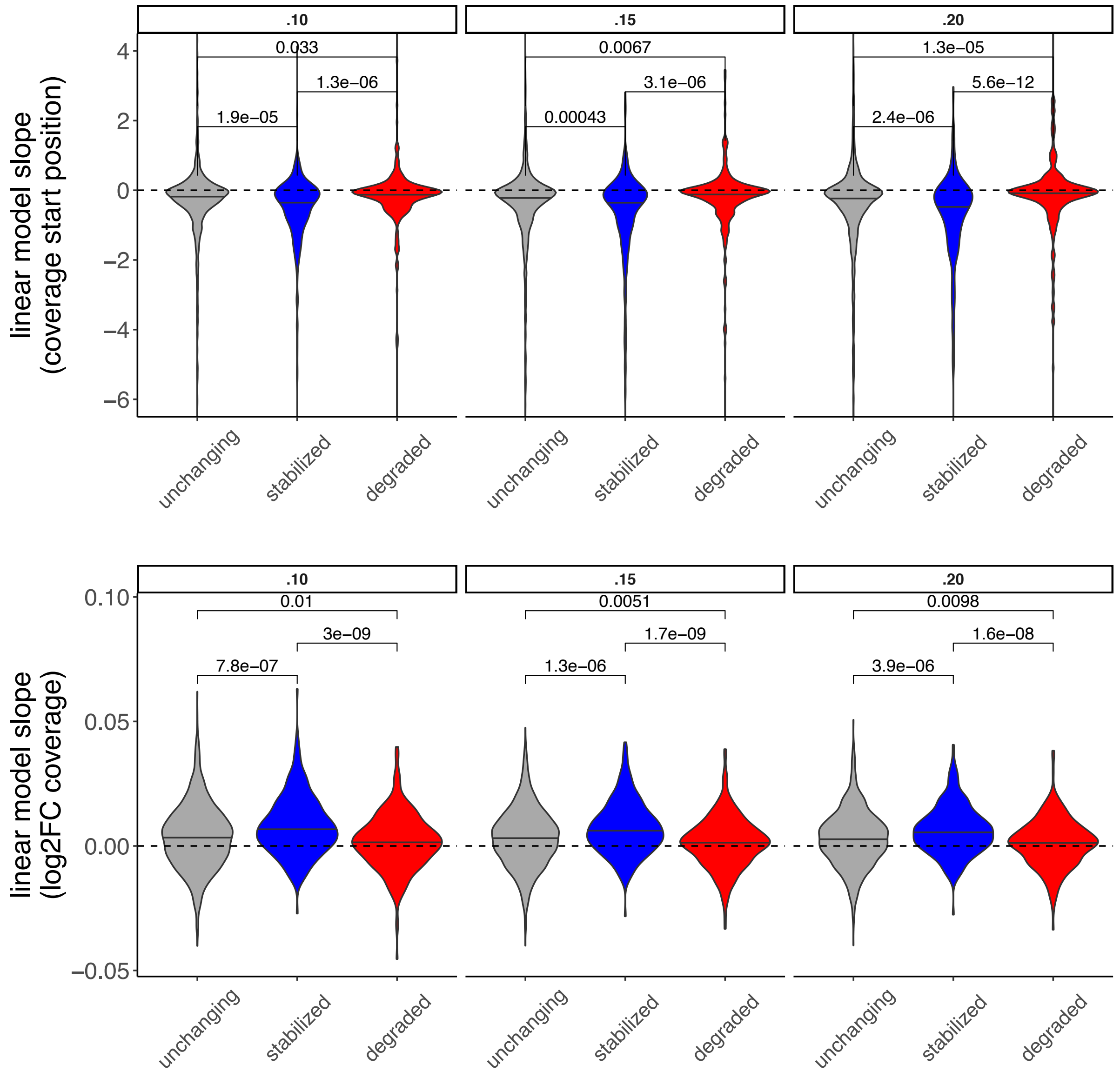


**Supplementary Figure 14**. Coverage differences between mRNAs are similar using different 5' cutoffs.

Differences between RNA-seq coverage in stabilized, unchanging and degraded mRNAs when using different cutoffs to define coverage starting points. Same strategy as shown in Figure 5A.

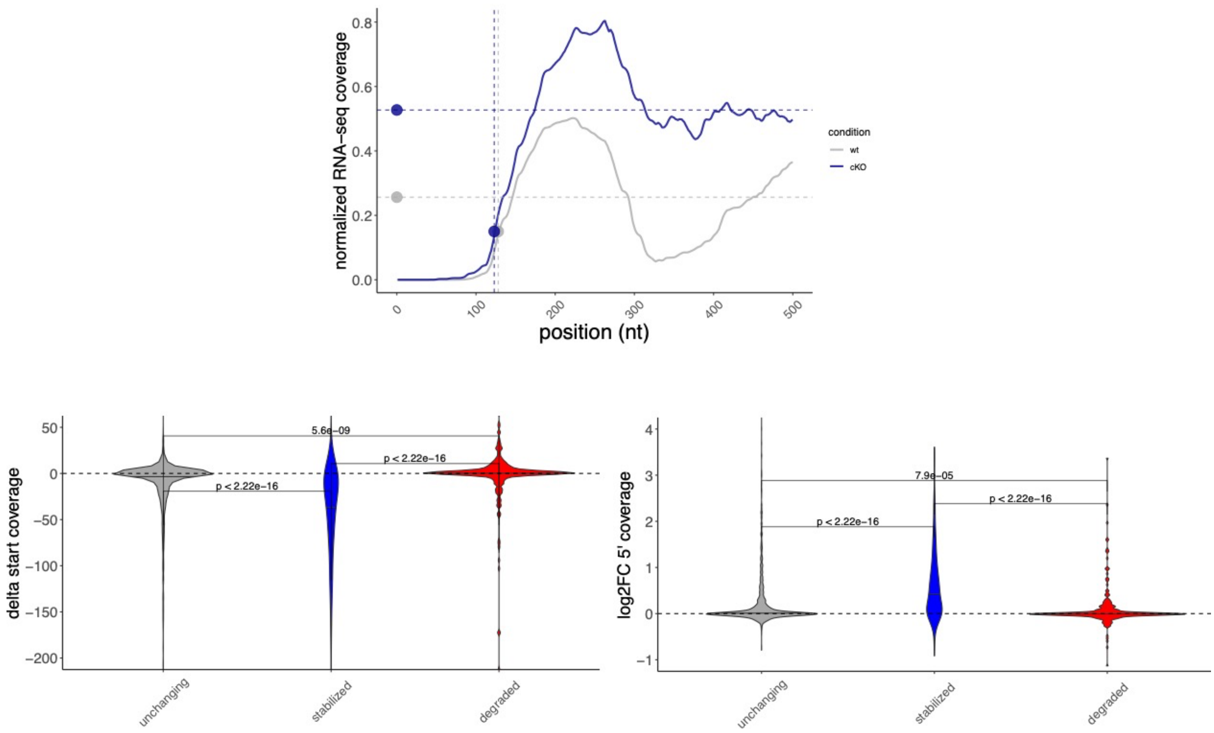


**Supplementary Figure 15**. Changes in RNA-seq coverage values and starting positions in the *Ddx3x* cKO mouse.

Exemplified strategy using an mRNA from the *Ctxn1* gene. Differences in coverage values (log2FC) and starting points are extracted and compared across regulated mRNAs (Methods). Coverage values were 0-1 normalized for each dataset.

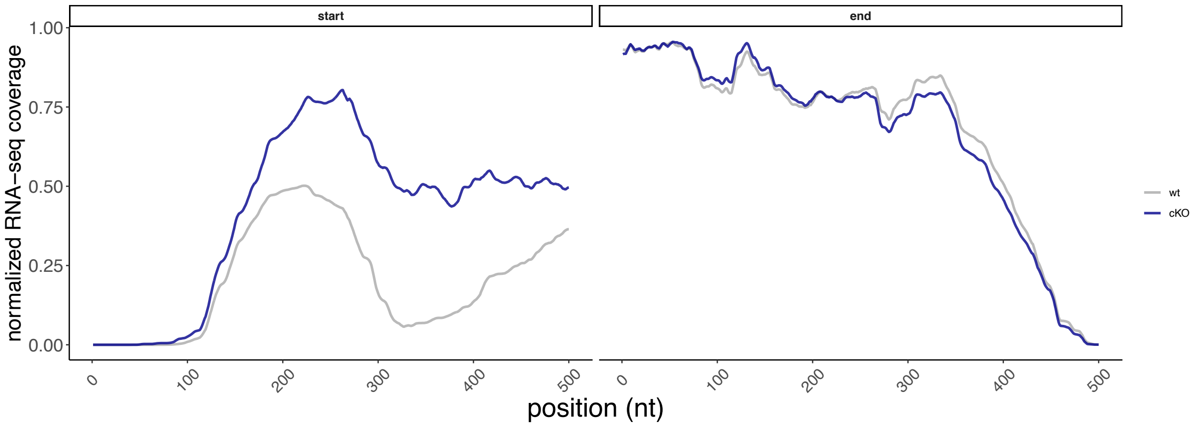


**Supplementary Figure 16**. Example of RNA-seq coverage changes in the *Ddx3x* cKO mouse.

RNA-seq coverage tracks around 5’ and 3’ ends of the stabilized mRNA from the *Ctxn1* gene. Coverage values were 0-1 normalized for each dataset.
